# Extracellular Vesicles Hijack the Autophagic Pathway to Induce Tau Accumulation in Endolysosomes

**DOI:** 10.1101/2020.05.27.118323

**Authors:** Giona Pedrioli, Marialuisa Barberis, Maurizio Molinari, Diego Morone, Stéphanie Papin, Paolo Paganetti

**Affiliations:** Neurodegeneration Research Group, Laboratory for Biomedical Neurosciences, Neurocenter of Southern Switzerland, Ente Ospedaliero Cantonale, Torricella-Taverne, CH-6807, Switzerland; Member of the International PhD Program of the Biozentrum, University of Basel, Basel, CH-4056, Switzerland; Università degli Studi dell’Insubria, Varese, I-21100, Italy; Protein Folding and Quality Control, Institute for Research in Biomedicine, Bellinzona, CH-6500, Switzerland; Imaging Facility, Institute for Research in Biomedicine, Bellinzona, CH-6500, Switzerland; Faculty of Biomedical Sciences, Università della Svizzera italiana, Lugano, CH-6900, Switzerland

**Keywords:** Autophagy, Endolysosome, Extracellular vesicles, Neurodegeneration, Tau

## Abstract

Clinical progression of tauopathies is reflected by the transcellular propagation of pathogenic Tau seeds with the possible involvement of extracellular vesicles as transport vectors. However, the mechanism regulating extracellular vesicle cargo delivery to recipient cells is poorly understood. We established a cell model for investigating extracellular vesicle-delivery of membranes and proteins. In this model, extracellular vesicles are readily internalized and accumulate in endolysosomes. For the first time, we show that in this acidic compartment of recipient cells, extracellular vesicle-delivered Tau seeds cause the accumulation and abnormal folding of normal Tau by a process that requires the participation of autophagy. Endolysomes represent thus a cross-road where Tau seeds released from extracellular vesicles propagate on cellular Tau on its route for autophagy-mediated degradation, ultimately driving its accumulation, endolysosomal stress and cytotoxicity. Whilst, autophagy stimulation is considered as a viable solution to protect neurons from harmful cytosolic protein inclusions, our data suggest that this approach may favour the aberrant accumulation of neurodegeneration-associated proteins induced by exogenous pathogenic protein forms, with possible implications in the spreading of the disease.

## Introduction

Tauopathies are a group of progressive neurodegenerative diseases characterized by intracellular fibrillar Tau protein species ultimately deposited in neurofibrillary tangles as the pathological disorder hallmark^1,2^. Tangles start to appear in vulnerable disease-specific brain regions and then gradually spread along the neuronal connectivities invading the whole brain. Indeed, the temporal and spatial tangle distribution correlates well with the extent of neuronal loss and, importantly, with the clinical disease stages^3^. Based on this, most aging-associated neurodegenerative disorders are described as proteinopathies (or misfolded protein disorders) characterized by a gain-of-toxic function linked to impaired protein homeostasis, protein deposition and proteotoxicity^4^.

Disease progression and brain propagation of pathogenic protein species involve a prion-like course where fibrillar Tau species are able of self-propagate through a still largely unclarified protein-to-protein mechanism, which needs to involve a cell-to-cell transmission^5,6^. Different means of communication account for transcellular transport of macromolecules. Among these, extracellular vesicles (EVs), which function as paracrine vectors, are emerging as deeply involved in pathological disease propagation^7^. EVs contribute to cancer^8^, inflammation disorders^9^ and also to neurodegeneration where EVs carrying replication-competent particles contribute to disease progression^10,11^.

EVs-mediated communication relies on transferring or displaying their cargo from a donor cell to a recipient cell^12^. RNAs are among the most studied EVs-cargo macromolecules. In recipient cells, EVs-mRNA are translated into protein effectors or EVs-miRNA regulates gene expression^9,13–15^. However, the evidence for a biological function of proteins transported by EVs remains sparse^16^, especially concerning the molecular mechanisms regulating EVs-cargo delivery in recipient cells. Direct fusion with the plasma membrane represents the simplest theoretical route of delivery^17^. However, EVs exploit the endocytic pathway as the main route for cell internalization^18–20^. This resembles the fate of (macroscopic) nutrients that after internalization reach endolysosomes (ELs) for degradation and recycling. ELs are acidic cellular organelles whose main function is the degradation of both intracellular and extracellular material, thus representing a major crossroad for two major degradative pathways, respectively endocytosis and autophagy^21^. Macroautophagy is a regulated degradation pathway for organelles and long-living cytosolic proteins and aggregates that bypass the ubiquitin-proteasome system^22,23^. Macroautophagy is characterized by the formation of double-bilayer autophagosomes that act as cargo delivery vectors for intracellular material to ELs^24,25^. Not surprisingly, ELs are key players in regulating cellular proteostasis and age-associated ELs impairment is a pronounced feature of neurodegenerative diseases, linked to cellular stress and cell death^26,27^.

Here, we report that ELs represent the subcellular site where EVs-delivered profibrillar Tau meets and initiate the accumulation of an intracellular Tau pool on route for autophagy-mediated degradation. Thus, our data propose that ELs represent a critical subcellular location for EVs cargo delivery with disease relevance. Indeed, we show that in cells internalizing profibrillar Tau transported by EVs, Tau accumulated within ELs causes a cell stress response and cytotoxicity. ELs-Tau also acquires a conformation and phosphorylation also observed when deposited in brain tangles.

## Materials and Methods

### Cell culturing

All cells were routinely grown at 37°C with saturated humidity and 5% CO_2_ in Dulbecco’s Modified Eagle Medium (DMEM, Gibco, 61965-059) supplemented with 1% non-essential amino acids (Gibco, 11140035), 1% penicillin-streptomycin (Gibco, 15140122), 10% fetal bovine serum (FBS, Gibco, 10270106) and passaged at 90% confluency. Inducible mouse multipotent neural progenitor C17.2 cells (ECACC 07062902) were generated with the Flp-In T-Rex tetracycline-inducible cell system according to the manufacturer instructions (Invitrogen, K650001) and maintained in the presence of 150 μg/mL hygromycin B (Invitrogen, 10687010) and 15 μg/ml blasticidin S (Gibco, A1113903). Gene expression was routinely induced in the presence of 60 ng/mL tetracycline. To induce autophagy, cells were washed three times with PBS and then incubated at 37°C for 4 h in Hanks’ Balanced Salt Solution (Gibco, 14025-050).

### Plasmids

Addgene plasmids used were: pCMV-lyso-pHluorin (RRID:Addgene_70113, gift from C. Rosenmund^28^), pLAMP1-mCherry (RRID:Addgene_45147, gift from A. Palmer^29^), mCherry-CD9-10 (RRID:Addgene_55013, gift from M. Davidson), mCh-Rab7A and mCh-Rab5 (RRID:Addgene_61804, RRID:Addgene_49201, gift from G. Voeltz^30,31^), pLV-CMV-LoxP-DsRed-LoxP-eGFP and pcDNA3.1-CMV-CFP;UBC-Cre25nt (RRID:Addgene_65726, RRID:Addgene_65727, gift from J. van Rheenen^15^). GFP-CD63 was obtained with sequential subcloning of the CD63 cDNA in the vector pCDNA5/FRT/TO followed by in-frame 5’-insertion of the GFP cDNA with HindIII/BlpI. GFP-Tau_MBD_ was generated by replacing the CD63 cDNA with that encoding for Tau_MBD_ with BamHI/XhoI. Tau_441_-mCherry was obtained with sequential subcloning of the Tau_441_-mCherry cDNA in the vector pCDNA5/FRT/TO with BamHI/XhoI. For the split GFP, Tau_441_-mCherry and Tau_MBD_ cDNAs were subcloned in parental plasmids as described previously^32^. Plasmid transfections were performed the day after cell plating with Lipofectamine 3000 (Invitrogen, L-3000-008), following the manufacturer’s instructions.

### Co-culture paradigms

For the analysis of transcellular transport of lipid membranes, cells stably integrated with the LoxP-DsRed-LoxP-eGFP cassettes were cultured with DiO or DiD labelled cells for 72 h in a well of a 8-well chamber (ibidi, 80826). For all other co-culture experiments DiO or DiD cells were replaced with cells with inducible expression of the protein product of interest. Treatment with 25 nM bafilomycin A1 (Sigma, B1793-2UG) lasted for 4 h and that with 33 μM dynasore (Sigma, D7693) for 2 hr. Analysis of the co-cultures were performed in-live by laser confocal microscopy, or with a benchtop flow cytometer (Beckman, CytoFLEX) after collecting cells with TrypLE™ (Gibco, 12604021) and resuspension of the cells in DMEM without phenol red (Gibco, 21063045) supplemented with 10% FBS.

### EVs enrichment

Conditioned serum-free medium for EVs isolation was routinely obtained after an incubation on cells for 72 hr. DiO-labelled, DiD-labelled of inducible C17.2 cells were first grown in 10 cm petri dishes (Corning, 353003) to 80% confluency in complete medium. Transiently transfected cells were washed twice with PBS at 4 h post-transfection before switching to the serum-free conditions. EVs were isolated from conditioned media by established procedures^33^. In short, dead floating cells and cell debris were removed by centrifugation at 1’000 g for 20 min at 4°C. EVs were concentrated with 100 kDa-cutoff filters (Sigma, UFC910096) at 3’000 g for 15 min at 4°C. The concentrated suspension was first cleared from large membrane vesicles by centrifugation at 10’000 g for 20 min at 4°C for 20 min, followed by serial centrifugation at 100’000 g for 120 min at 10 °C (Beckman fixed-angle TL110 rotor; Beckman Optima Max-TL ultracentrifuge). The so obtained P100 pellets, corresponded to the enriched EVs fraction, were re-suspended in 10 μL/10 cm dish in ice-cold PBS. Size and concentration of EVs was determined through nanoparticle tracking analysis (NanoSight, LM10) after each EVs preparation with at least three repeated quantifications for each preparation. For routine cell uptake experiments, cells were incubated with 10^9^ freshly-enriched EVs for 16 h, or differently explained in figure legends, on 15’000 cells/cm^2^ in an 8-well chamber.

### Size exclusion chromatography

For EVs-purification by size exclusion chromatography (SEC), P100 (pellet) or S100 (supernatant) obtained from the final 100’000 g centrifugation were separated on a qEVoriginal/70nm column (Izon Science, SP1) topped with 14 mL PBS. 500 μL fractions were collected and fraction 7 to 24 were characterized by nanoparticle tracking analysis and for protein concentration. Fractions were then concentrated on 30 kDa-cutoff filters (Sigma, UFC803024) and stored frozen at −20°C.

### Staining of cells

For immune-fluorescence microscopy, cells routinely grown on poly-D-lysine coated 8-well chambers, were fixed in freshly diluted 4% formaldehyde (Sigma-Aldrich, F1635) for 15 min at RT, washed twice with PBS supplemented with 100 mM glycine for 5 min and once with PBS. Cells were permeabilized in 0.05% saponin (Sigma-Aldrich, 84510) or 0.1% triton (Sigma-Aldrich, X100) in PBS for 10 min and blocked with 5% normal goat serum (Biowest, S2000-500) in PBS for 30 min before incubation with primary antibodies for 1 h at RT in the permeabilization buffer supplemented with 0.5% normal goat serum: TFE3 (0.6 μg/ml; Sigma-Aldrich, HPA023881), LAMP1 (1/50 conditioned medium of hybridoma cells; DSHB Hybridoma Product 1D4B was deposited by J.T. August), MC1 (1/100 conditioned medium of hybridoma cells; kindly provided by Peter Davies), AT8 (0.2 μg/ml, Thermo Fischer Scientific, MN1020). Detection was performed with secondary antibodies (2 μg/mL at RT in the dark for 1 h) α-mouse IgG AlexaFluor™ 350 (Thermo Fischer Scientific, A21049), α-rabbit IgG AlexaFluor™ 647 (Thermo Fischer Scientific, A21245), α-rabbit IgG AlexaFluor™ 488 (Thermo Fischer Scientific, A11034) and α-rat IgG AlexaFluor™ 647 (Thermo Fischer Scientific, A21247). Nuclei were counterstained with 0.5 μg/mL DAPI (Sigma-Aldrich, D9542).

For all the experiments requiring fluorescent membrane labelling, cells were incubated in 1 mL solution containing 5 μL Vybrant DiO (Molecular Probes, V22886) or Vybrant DiD (Molecular Probes, V22887) at 37°C for 10 min, followed by extensive washing in excess PBS. For bulk endocytosis studies, cells previously seeded overnight were incubated for 4 h with 10 μM Alexa Fluor™ 594-dextran (Invitrogen, D22913) and analysed in live by laser confocal microscopy.

For the identification of ELs acidic compartments, cells were incubated with 75 nM LysoTracker Red DND-99 or Deep Red (Thermo Fischer Scientific, L7528 and L12492) in complete medium at 37 °C for 30 min, washed with an excess of PBS and either imaged live by laser confocal microscopy or detached with trypsin and resuspended in DMEM without phenol red added with 10% FBS for cytometric analysis.

### Western blots

For biochemical analysis by western blot, cells grown in 10 cm plates were washed once in PBS, directly lysed in SDS-PAGE sample buffer (1.5% SDS, 8.3% glycerol, 0.005% bromophenol blue, 1.6% β-mercaptoethanol and 62.5 mM Tris pH 6.8) and boiled at 96°C for 10 min. EVs preparations were also lysed in SDS-PAGE sample buffer. Equal volume of cell (representing 1/20 of the total lysate) and EVs lysates (representing total lysate) were resolved by SDS-PAGE and transferred to PVDF membranes (BioRad, 162-0177), blocked in Odyssey Blocking Buffer (LI-COR, 927-50000) and visualized with the infrared western blot technology on Odissey CLx device (LI-COR). Primary antibodies were incubated for 1 h at 37°C and were specific for ALIX (1 μg/mL; abcam, ab117600), TSG101 (1 μg/mL; abcam, ab30871), TOMM20 (0.8 μg/mL; abcam, ab78547), GFP and pHluorin (5 μg/mL; abcam, ab290), EEA1 (1/500; Thermo Fischer Scientific, MA5-14794), LC3B (1 μg/mL; Novus Biologicals, NB600-1384), TAU13 (0.2 μg/mL; Santa Cruz, sc-21796), GAPDH (0.18 ug/mL; abcam, ab181602), β-actin (0.10 μg/mL; Sigma-Aldrich, A1978) and 2.3 μg/mL β1^34^. Secondary antibodies were incubated for 1 h at 37°C and were α-rabbit IgG coupled to IRDye 800CW (LI-COR) and α-mouse IgG coupled to IRDye 680RD (LI-COR).

### Microscopic image analysis

Fluorescence images were acquired with either a confocal microscope (Nikon C2) or Lionheart FX automated microscope (Biotek). Image analysis and processing of the raw data were performed with the Gen5 (BioTek) or Fiji/ImageJ (1.51 g or later) software. For flow cytometry analysis, cells fluorescence was acquired on CytoFLEX analytical device (Beckman Coulter). The ImageJ “JACoP” plugin was used to quantify Pearson’s correlation coefficient (PCC) and Manders’ overlapping coefficient (MOC, M1 & M2) for dual-colors co-localization. Alternatively, when a cellular mask was required (i.e. in transient transfected cells), the imageJ “Coloc2” plugin was used to quantify both PCC and MOC.

For Tau-puncta quantification, C17.2 cells expressing Tau_441_ and incubated in culture medium supplemented with vehicle (PBS), P-EVs or Tau_MBD_-EVs were imaged by laser confocal microscopy. Quantification of the percentage of cells with a Tau_-_puncta phenotype was normalized with the total number of DAPI-positive nuclei. Single Tau-puncta were manually identified and analyzed with the ImageJ software.

For nuclear TFE3 quantification, cells were fixed, stained for TFE3 and imaged by laser confocal microscopy. A DAPI-nuclear mask was applied to determine TFE3 fluorescence intensity within the nucleus and outside this mask. Quantification of the percentage of cells with a nuclear TFE3 phenotype was then determined applying an arbitrary threshold to the ratio between the two resulted masks.

### Automated ELs analysis

For the identification of ELs, cells were stained live with LysoTracker. Images acquired at confocal microscopy were analysed with Fiji/ImageJ. Automatic segmentation of LysoTracker positive organelles was performed with a Weka machine learning approach. Training set comprised three images in which LysoTracker positive ELs and background regions were manually annotated. After balancing classes, Weka was trained with Gaussian Blur (sigma 1-16), Sobel filter, Hessian, Difference of gaussians, Membrane projections and Lipschitz features. We then applied watershed filter to the output mask and filtered objects smaller than 0.2 μm^2^ to generate a set of ROIs for LysoTracker positive ELs. Manual annotation of the cells of interest on each image, based on Tau_441_, were then used to create ROIs for the cytoplasm, by subtracting the combined ROIs of LysoTracker positive ELs from the ROI of the cell. ELs size and mean intensity were then quantified on biFC and Tau_441_ channels using this set of ROIs.

### Cell toxicity assays

LDH assay (Pierce LDH Cytotoxicity Assay Kit; 88954, ThermoFisher Scientific) was performed following the manufacturer’s instructions. In short, conditioned medium from cells subjected to EVs treatments was collected, processed and analyzed by measuring the absorbance at 490 nm and 680 nm in an Infinite M Plex (Tecan).

The nuclear area and the cell area were analyzed as a proxy for early cell toxicity events. Cells treated with EVs were stained with Hoechst at 37°C for 5 min, washed with PBS, fixed, imaged at Lionheart microscope and analyzed with the Gen5 software. The mean nuclear area was analyzed based on a Hoechst positive nuclear mask. The mean cell area was quantified measuring a cell mask based on Tau_441_.

### Statistics and reproducibility

All the experiment reported were repeated at least in three independent experiments if not otherwise explained in the legend of the figure. Differences between two means were assessed by unpaired non-parametric Mann-Whitney test, whereas differences between more than two means were tested by non-parametric Kruskal-Wallis one-way ANOVA, alpha = 0.05, if not otherwise indicated in figure legends. A P value < 0.05 was used to assess significances. Statistical analysis and graphs were generated with Prism (GraphPad software version 8).

### Data availability

All raw data supporting the findings of the study are available from the authors upon request.

## Results

### A cell paradigm to investigate EVs transport and internalization

We first validated our cell system as a cellular paradigm granting transcellular exchange of EVs whilst allowing to dissect the process of aberrant cell-to-cell propagation of Tau. We opted for myc-immortalized mouse cerebellar C17.2 cells, a cell line generated for studying neural progenitor cells^35^ and seed-induced aggregation of Tau^6,36^.

In order to evaluate whether C17.2 cells were able to exchange EVs, we first co-cultured two cell populations, one labelled with DiO, a lipophilic fluorescent tracer (emission maximum Em_max_ = 501 nm) that intercalates in cellular membranes and one expressing the fluorescent protein DsRed (Em_max_ = 583 nm). Cells were co-cultured at a 5:1 ratio (DiO:DsRed) for 72 h (Fig 1A). Analysis by liquid flow-cytometry revealed a substantial fraction of co-cultured cells positive for both markers when compared to parental cells or to single cultures of DiO and DsRed cells (Fig 1B & C). Similar results were obtained when DiO was replaced with DiD (Em_max_ = 665 nm) under the same assay conditions (Supplementary Fig 1A). Notably, the predominant population behaving as recipient cells was the DsRed population since it acquired the double fluorescent properties. In fact, the proportion of DsRed cells in the co-culture for which a single tracer was still detectable was only ~17% of the total DsRed population (Supplementary Fig 1B). Preferential EVs transport of the fluorescent tracers DiO or DiD possibly relied on their reduced size when compared to the ~100-time larger tetrameric DsRed or on their association to membranes. Analysis by confocal laser scanning microscopy of the co-culture showed that DsRed-expressing recipient cells internalized DiO-stained donor cell-membranes in a dot-like patter that was evocative of the endocytic system (Fig 1D). Also, the orthogonal reconstruction of serial confocal planes confirmed that DiO-membranes were internalized and located within the recipient cell (Fig 1D). The DiO/DsRed or DiD/DsRed co-culture assay enabled to monitor unidirectional membrane transport with DsRed-fluorescence identifying the recipient cells that internalized membranes originating from DiO or DiD-labelled donor cells. In order to directly assess the involvement of EVs, these were enriched starting from serum-free conditioned medium of donor cells by a standard protocol based on serial centrifugations^37^. The EVs fraction was first characterized for size and concentration by nanoparticle tracking analysis^38^. The more frequent EVs population presented a diameter of 177 ± 13 nm (Fig 1E). Western blot analysis of EVs demonstrated the relative enrichment of the EVs markers Alix and TSG101 when compared to donor cell lysates (Fig 1F). In contrast, markers of potentially contaminating membranes, early endosomes (EEA1) and mitochondria (TOMM20), displayed an opposite behaviour (Fig 1F). EVs isolated from the conditioned medium of DiD-labelled donor cells were then incubated on DsRed-expressing recipient C17.2 cells (Fig 1G). This established a dose-dependent accumulation of DiD-fluorescence, which gradually increased over the 0 to 16 h incubation time (Fig 1H).

**Figure 1.**
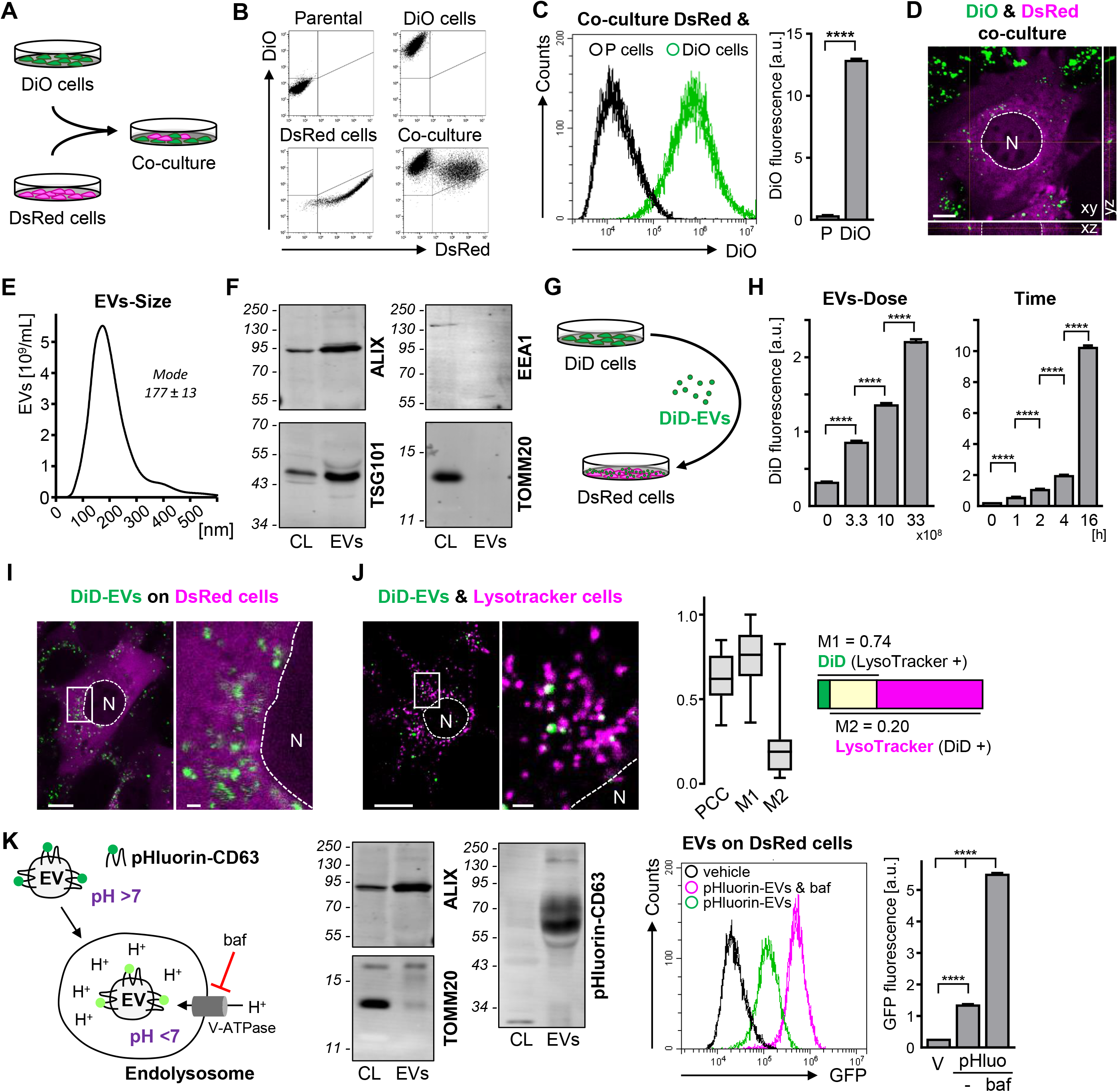
A cell paradigm to investigate EVs transport and internalization. A Scheme of co-culture assay for monitor transcellular transport of macromolecules (DiO in green, DsRed in magenta). B 2D scatter plots of single cell DsRed and DiO fluorescence by flow-cytofluorimetry for the indicated cell populations (n ~5 ×10^3^ cells/condition). C 1D plots of DiO-fluorescence intensity for single DsRed cells co-cultured with parental cells (P cells) or DiO-labelled cells (DiO cells), the data are pooled from five flow-cytofluorimetry analyses (n ~5 ×10^3^ cells/replicate). The histogram shows the DiO fluorescence intensity determined for DsRed cells co-cultured as indicated (mean ± s.e.m. of six biological replicates with 48.6 ×10^3^, 57.1 ×10^3^ cells respectively). **** P<10^−15^, unpaired non-parametric Mann-Whitney test. D Fluorescent confocal image of a DiO & DsRed co-culture (DiO in green, DsRed in magenta), the image includes an orthogonal reconstruction of 14 focal z-planes demonstrating internalized DiO-stained membranes in a DsRed cell. Scale bar: 10 μm. E Size distribution of a representative EVs preparation determined by nanoparticle tracking analysis. The size of the most represented EVs population is given (mode ± s.d. of three technical replicates, n = 1.1 ×10^3^, 0.9 ×10^3^, 0.9 ×10^3^ particles respectively). F Western blots of donor cell lysates (CL) and EVs lysates (EVs) for the EVs markers Alix and TSG101, or for potentially contaminating endosomes (EEA1) or mitochondria (TOMM20). Molecular weight markers are given on the left. G Scheme of the DiD-EVs assay on acceptor DsRed cells (DiD in green, DsRed in magenta). H DiD-fluorescence intensity in single DsRed cells incubated with the indicated amount of DiD-EVs for 2 h (mean ± s.e.m. of three biological replicates with 8.6 ×10^3^, 7.2 ×10^3^, 8.0 ×10^3^, 7.9 ×10^3^ cells respectively) or with 1 ×10^9^ DiD-EVs for the indicated times (mean ± s.e.m. of three biological replicates with 8.3 ×10^3^, 11.0 ×10^3^, 10.0 ×10^3^, 9.1 ×10^3^, 9.2 ×10^3^ cells respectively). ****P<10^−15^ non-parametric Kruskal-Wallis one-way ANOVA, alpha = 0.05. I Fluorescent confocal image of DsRed cells incubated for 4 h with 1 ×10^9^ DiD-EVs (DiD in green, DsRed in magenta, N = nucleus). Scale bar: 10 μm (1 μm for the enlarged inset). J Fluorescent confocal image of C17.2 cells incubated for 16 h with 1 ×10^9^ DiD-EVs (in green) and counterstained with 75 nM LysoTracker (in magenta); co-emission of the two fluorophores results in a white colour and indicate localisation of EVs within acidic organelles. Scale bar: 10 μm (1 μm for the enlarged inset). Shown are also Pearson’s correlation coefficient (PCC) and Manders’ overlapping coefficient (M1 & M2) for 33 images from a representative experiment: 1-99 percentiles (whiskers), 25-75 percentiles (box), median (horizontal bar within the box). The majority of DiD puncta localizes with the relative minority of LysoTracker-organelles. K Scheme of pHluorin fluorescence quenching by the acidic ELs environment upon pHluorin-CD63-EVs internalization. Maximal pHluorin fluorescence at neutral pH is shown in dark green and the pH<7 quenched emission in light green. Acidification of ELs occurs though active proton (H^+^) transport by the proton pump (V-ATPase) inhibited by bafilomycin (baf). L Western blots of donor cell lysates (CL) and EVs lysates (EVs) for the EVs-markers Alix and TSG101, or with a GFP-antibody detecting pHluorin-CD63 that is highly EVs enriched. Molecular weight markers are given on the left. M 1D plots of single cell pHluorin-fluorescence intensity by flow-cytofluorimetry for cells incubated with 1 ×10^9^ pHluorin-CD63-EVs (pHluorin-EVs) for 16 h in the absence (in green) or presence (in magenta) of 25 nM bafilomycin-A1 co-supplemented for the last 4 h (pHluorin-EVs & baf;) or without EVs (vehicle, in black). The histogram shows GFP-fluorescence intensity for the given treatments (mean ± s.e.m. of three biological replicates with 12.6 ×10^3^, 9.5 ×10^3^, 12.1 ×10^3^ cells respectively), ****P<10^−15^, non-parametric Kruskal-Wallis one-way ANOVA, alpha = 0.05.

Ectopic expression of the ELs markers Rab5, Rab7, CD9 and LAMP1 fused to mCherry (emission maximum at 610 nm) in acceptor cells established that donor membranes accumulated in ELs of recipient cells at 16 h incubation (Supplementary Fig 2A). This process may have occurred at least in part by a bulk-flow course similar to that used for internalization of 10 μM TRITC-labelled dextran supplied to the culture medium in combination to EVs (Supplementary Fig 2B). In the presence of 33 μM dynasore, a GTPase inhibitor that also interferes with lipid raft dynamics and clathrin-mediated endocytic pit formation^39^, mean DiD-fluorescence intensity was reduced by ~55% in recipient cells when compared to untreated cells (Supplementary Fig 2C). Confocal laser scanning microscopy analysis of recipient cells treated with DiD-EVs visualized a distribution patter of internalized EVs similar to that observed in the co-culture experiment (Fig 1I; compare to Fig 1D). Accumulation of internalized EVs in an acidic compartment of recipient cells was further supported by the co-localization of DiD-EVs with the tracer LysoTracker (Fig 1J). Determination of the Manders’ overlapping coefficient (MOC) showed that ~74% of the DiD-positive organelles were LysoTracker-positive ELs (M1), whereas ~20% of LysoTracker-positive ELs contained EVs-DiD (M2) (Fig 1J). Moreover, EVs loaded with the pH-sensitive biofluorescent protein pHluorin^40^ (Em_max_ = 509 nm) in recipient cells that were treated with a proton-pump inhibitor, displayed increased fluorescence intensity when compared to untreated conditions (Fig 1K). In conclusion we found that C17.2 cells represented an adequate model to analyse transcellular exchange of EVs and that EVs are internalized by an endocytic process in recipient cells as reported previously^41^.

**Figure 2.**
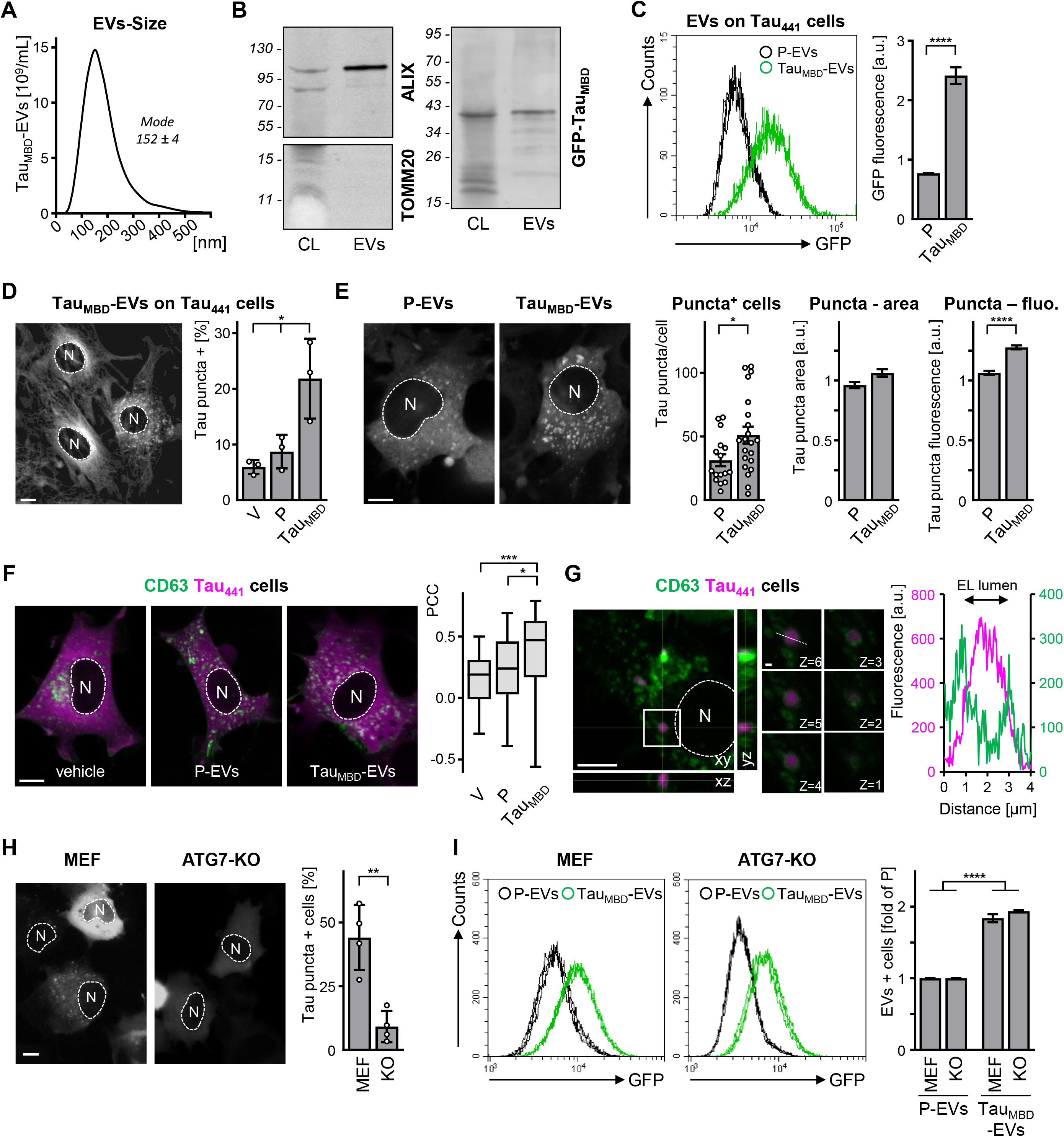
EVs transported Tau_MBD_ induces cellular Tau_441_ accumulation in ELs via autophagy. A Size distribution of a representative Tau_MBD_-EVs preparation determined by nanoparticle tracking analysis. The size of the most represented EVs population is given (mode ± s.d. of three technical replicates with 1.6 ×10^3^, 1.5 ×10^3^, 1.6 ×10^3^ particles respectively). B Western blots of donor cell lysates (CL) and EVs lysates (EVs) for the EVs markers Alix and TSG101, or with a GFP-antibody detecting GFP-Tau_MBD_. Molecular weight markers are given on the left. C 1D plots of single cell GFP-fluorescence intensity by flow-cytofluorimetry for Tau_441_ cells incubated with 1 ×10^9^ parental EVs (P-EVs, in black) or GFP-Tau_MBD_-EVs (Tau_MBD_, in green) for 24 h, data are pooled from three biological replicates (~4.6 ×10^3^ cells/replicate). The histogram shows GFP-fluorescence intensity for the given treatments (mean ± s.e.m. of three biological replicates with 14.9 ×10^3^, 15.9 ×10^3^ cells respectively), ****P<10^−15^, unpaired non-parametric Mann-Whitney test. D Fluorescent image of Tau_441_ cells incubated for 24 h with 1 ×10^9^ Tau_MBD_-EVs (Tau_441_ in grey, N = nucleus). Scale bar: 10 μm. The histogram shows % of cells with a Tau-puncta phenotype for the given treatments: vehicle (V), parental-EVs (P) or Tau_MBD_-EVs (Tau_MBD_) (mean ± s.d. of three biological replicates with 2.2 ×10^3^, 4.0 ×10^3^, 4.7 ×10^3^ cells respectively). *P<0.0288, ordinary one-way ANOVA with Tukey post-hoc test, alpha = 0.05. E Fluorescent confocal image of Tau_441_ cells incubated for 24 h with either 1 ×10^9^ P-EVs (P) or 1 ×10^9^ Tau_MBD_-EVs (Tau_MBD_) (Tau_441_ in grey, N = nucleus). Scale bar: 10 μm. The histograms show the number of Tau-puncta per cell (mean ± s.d., n = 16, 21 cells respectively), area of puncta and Tau_441_ puncta fluorescence (mean ± s.e.m., n = 480, 1070 puncta respectively) for the given treatments. *P=0.0325, ****P<10^−15^, unpaired non-parametric Mann-Whitney test. F Fluorescent confocal image of cells expressing Tau_441_ and GFP-CD63 (CD63) incubated for 24 h with either vehicle (V), 1 ×10^9^ P-EVs (P) or 1 ×10^9^ Tau_MBD_-EVs (Tau_MBD_) (Tau_441_ in magenta, CD63 in green, N = nucleus), co-emission of the two fluorophores results in a white colour and indicate localisation of Tau_441_ within CD63 positive organelles. Scale bar: 10 μm. Shown is Pearson’s correlation coefficient (PCC) for 29, 31, 34 cells respectively from three biological replicates: 1-99 percentiles (whiskers), 25-75 percentiles (box), median (horizontal bar within the box). *P=0.0457, ***P=0.0005, non-parametric Kruskal-Wallis one-way ANOVA, alpha = 0.05. G Fluorescent confocal image of a cell expressing Tau_441_ and GFP-CD63 (CD63) incubated for 24 h with 1 ×10^9^ Tau_MBD_-EVs (Tau_MBD_) (Tau_441_ in magenta, CD63 in green, N = nucleus), the image includes an orthogonal reconstruction of 6 focal z-planes and z-planes series demonstrating a single enlarged Tau-puncta enclosed in CD63 positive membranes. Scale bar: 10 μm (1 μm for the enlarged inset). The scatter plot shows Tau_441_- and CD63-fluorescence from the dotted-line represented in z=6. Scale bar: 10 μm (1 μm for the enlarged inset). H Fluorescent images of MEF cells expressing Tau_441_ incubated for 24 h with 1 ×10^9^ Tau_MBD_-EVs (Tau_441_ in grey, N = nucleus). Scale bar: 10 μm. The histogram shows the % of cells with a Tau-puncta phenotype for the given recipient cells: parental MEF (MEF) or ATG7-KO MEF (KO) cells (mean ± s.d. of four biological replicates with 1.0 ×10^3^, 1.5 ×10^3^ cells respectively). *P=0.0026, two-tailed unpaired Student t-test. I 1D plots of single cell GFP-fluorescence intensity by flow-cytofluorimetry for Tau_441_ MEF cells (MEF and KO) incubated with 1 ×10^9^ parental EVs (P-EVs, black) or 1 ×10^9^ GFP-Tau_MBD_-EVs (Tau_MBD_, green) for 24 h, data are pooled from three biological replicates (58.9 ×10^3^, 59.8 ×10^3^, 59.0 ×10^3^, 59.4 ×10^3^ cells respectively). The histogram shows GFP-fluorescence intensity for the given treatments expressed as fold increase of the respective P-EVs treatment (mean ± s.e.m. of three biological replicates), ^ns^P<0.05, ****P<10^−15^, two-way Anova with Tukey post-hoc test, alpha = 0.05.

### EVs-mediated transport of proteins to ELs of recipient cells

In order to focus our attention to EVs-mediated transcellular transport of proteins, we next generated a cell line expressing a hybrid protein composed of GFP (Em_max_ = 510 nm) and CD63, a membrane-bound tetraspanin protein family member targeted to ELs and to EVs^42^. Co-culture of GFP-CD63 and DsRed cells at a 5:1 ratio led to the appearance of double-fluorescent cells when analysed by cytofluorimetry (Supplementary Fig 3A). Again suggesting preferential transport of GFP-CD63 when compared to DsRed, we noticed that most DsRed-cells acquired the GFP fluorescence. The relative fluorescent generated by GFP in double stained cells was lower to that observed when using the more concentrated and more brightly fluorescent DiO or DiD molecules. Orthogonal reconstruction by confocal microscopy showed internalized GFP-fluorescence with a dot-like distribution resembling that obtained with the DiO or DiD-fluorescent tracers (Supplementary Fig 3B, compare to Fig 1D and I). This GFP distribution pattern was clearly distinct from that observed in the GFP-CD63 expressing donor cells, which presented a more homogenous cell membrane GFP distribution in the absence of DsRed-fluorescence (Supplementary Fig 3B). The EVs-fraction isolated from the conditioned medium of donor cells displayed an enrichment of GFP-CD63 protein that was superior to that of endogenous Alix when compared to the respective amount detected by western blot in total cell lysates (Supplementary Fig 3C). GFP-CD63-EVs were internalized by DsRed recipient cells as shown by cytofluorimetry in cells incubated with either GFP-CD63- or parental-EVs (Supplementary Fig 3D). Internalized GFP-CD63-EVs co-localized with ectopic expressed LAMP1 (Supplementary Fig 3E). This cell assay enabled monitoring transcellular transport and internalization of EVs-cargo membrane-bound proteins.

**Figure 3.**
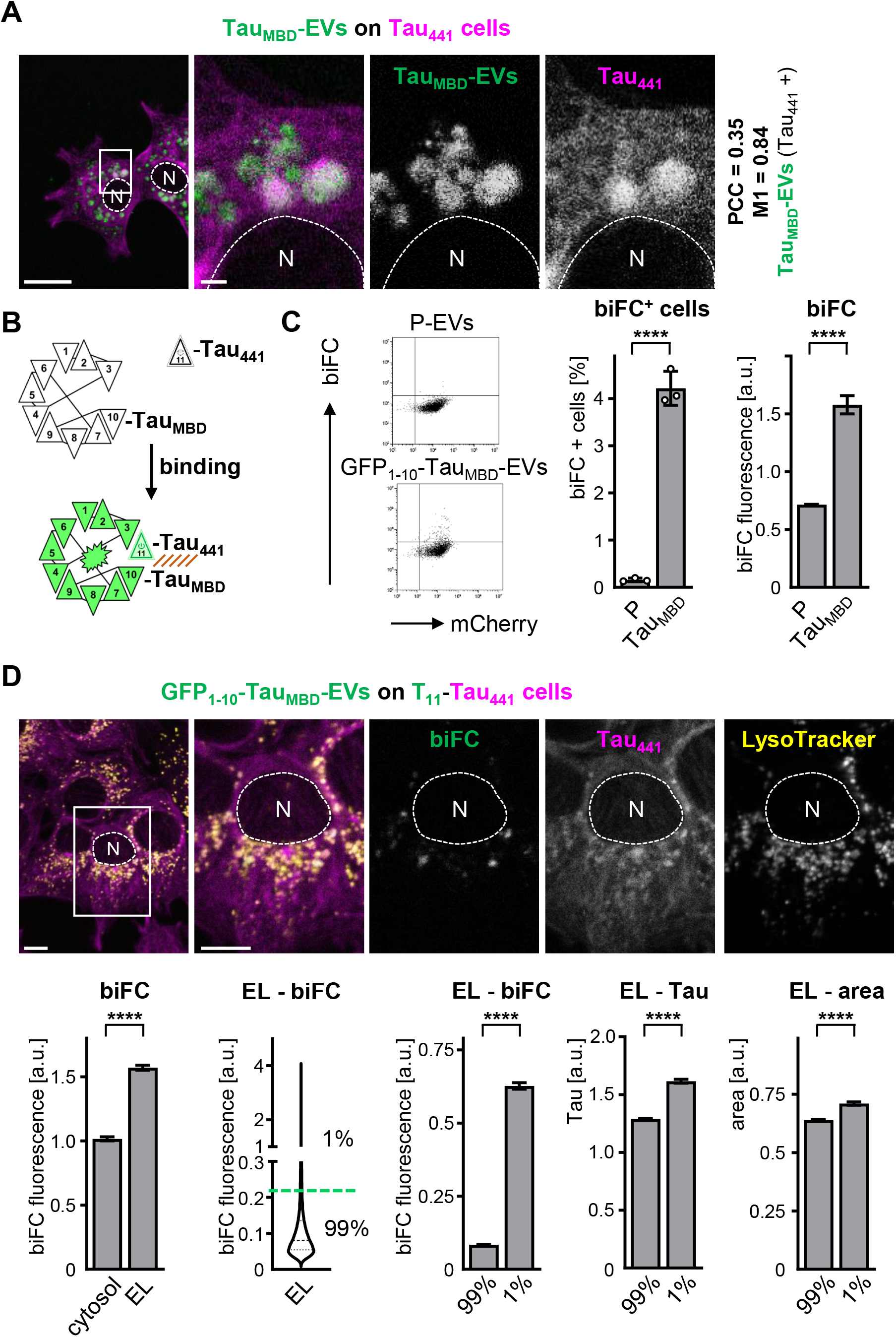
EVs transported Tau_MBD_ binds to recipient’s cells Tau_441_ in ELs. A Fluorescent confocal image of a Tau_441_ cell (in magenta) incubated for 72 h with 6 ×10^9^ GFP-Tau_MBD_-EVs (in green), co-emission of the two fluorophores results in a white colour and indicate localisation of GFP-Tau_MBD_-EVs within Tau-puncta. Scale bar: 10 μm (1 μm for the enlarged inset). B Scheme of the bimolecular fluorescence reconstitution (biFC) based on GFP. The first to tenth β-strands of GFP are fused to Tau_MBD_ and expressed in donor cells; the remaining eleventh β-strand of the GFP is fused to Tau_441_ and expressed in recipient cells. C 2D scatter plots of single cell Tau_441_ and biFC fluorescence by flow-cytofluorimetry for cells incubated for 72 h with either 6 ×10^9^ P-EVs or 6 ×10^9^ GFP_1-10_-Tau_MBD_-EVs (n = 1.2 ×10^3^, 0.8 ×10^3^ cells respectively). The histograms show the % of T_11_-Tau_441_ cells with biFC-fluorescence (mean ± s.d. of three biological replicates, ****P=4 ×10^−5^, two-tailed unpaired Student t-test) and biFC fluorescence intensity in Tau_441_ cells incubated with 6 ×10^9^ P-EVs (P) or 6 ×10^9^ GFP_1-10_-Tau_MBD_-EVs (Tau_MBD_) (mean ± s.e.m. of three biological replicates with 1.2 ×10^3^, 0.8 ×10^3^ cells respectively, ****P<10^−15^, unpaired non-parametric Mann-Whitney test). D Fluorescent confocal image of T_11_-Tau_441_ cells (in magenta) incubated for 72 h with 6 ×10^9^ GFP_1-10_-Tau_MBD_-EVs (biFC in green) and stained for LysoTracker (in yellow), co-emission of the three fluorophores results in a white colour and indicate localisation of biFC within LysoTracker positive Tau-puncta. Scale bar: 10 μm. The first histogram shows biFC-fluorescence intensity inside and outside the LysoTracker mask (mean ± s.e.m., n= 193 cells from three biological replicates, **** P<10^−15^, paired non-parametric Wilcoxon test). The violin plot shows the distribution of biFC-fluorescence in LysoTracker positive ELs (n= 2.7 ×10^3^ ELs), the green dotted-line represent the 1% outliers. The last three histograms show biFC-fluorescence, Tau_441_-fluorescence and area of LysoTracker positive ELs (mean ± s.m, n= 2.7 ×10^3^ ELs, ****P<10^−15^, unpaired non-parametric Mann-Whitney test).

### EVs transported profibrillar Tau induces Tau_441_ accumulation in ELs

Next, we investigated EVs-mediated transport of a protein lacking a transmembrane domain. For this, we utilized cells expressing a fibrillogenic fragment of Tau (Tau_MBD_) fused to GFP in order to isolate EVs with the mostly represented diameter of 152 ± 4 nm (Fig 2A). GFP-Tau_MBD_ was targeted to the EVs fraction less efficiently than GFP-CD63 and was detected at a relatively lower level than the endogenous EVs-marker Alix, but at a significantly higher level than the potential contaminant TOMM-20 (Fig 2B). In order to demonstrate that GFP-Tau_MBD_ was incorporated within EVs, we fist analysed by size-exclusion chromatography (SEC) the EVs-fraction, i.e. the pellet isolated by the last 100’000 g centrifugation (Supplementary Fig 4A). As expected pelleted EVs were recovered in the first SEC fractions representing the flow-through, whereas the main pool of soluble proteins present in the centrifugation supernatant were recovered in the following fractions representing the material with slower SEC migration (Supplementary Fig 4B). When comparing the EVs-pool to the soluble protein pool, GFP-Tau_MBD_ as well as endogenous Alix appeared enriched in EVs (Supplementary Fig 4C). Pretreatment of the EVs pool with the detergent Tx-100 inverted this behaviour (Supplementary Fig 4C). Furthermore, GFP-Tau_MBD_ co-isolated in the EVs-pool was largely protected from trypsin digestion in contrast to GFP-Tau_MBD_ recovered in the soluble protein pool (Supplementary Fig 4C). These data showed that the EVs-isolation procedure enriched for GFP-Tau_MBD_ targeted to the lumen of EVs. Consistent with this, we observed EVs-mediated transport of Tau_MBD_ with the determination of acquisition of GFP fluorescence in recipient cells by flow-cytofluorimetry (Fig 2C).

**Figure 4.**
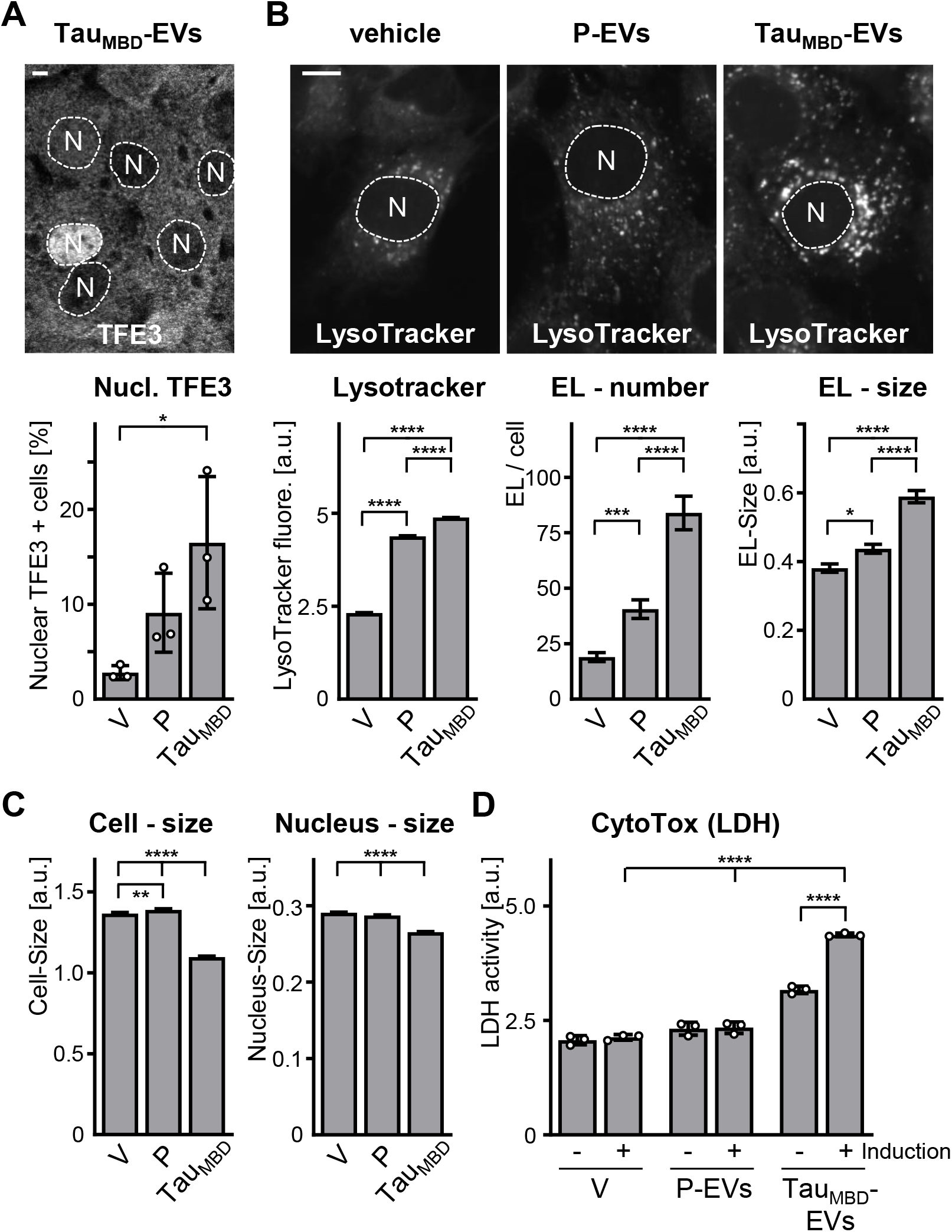
EVs-mediated Tau_441_ accumulation causes lysosomal stress and cytotoxicity. A Fluorescent confocal image of a Tau_441_ cell incubated for 24 h with either vehicle (V), 1 ×10^9^ parental-EVs (P) 1 ×10^9^ Tau_MBD_-EVs (Tau_MBD_) and stained for TFE3 (in grey, N = nucleus). The histogram shows the % of cells with a nuclear TFE3 localization for the given treatments (mean ± s.d. of three biological replicates with >1.2 ×10^3^ cells/condition). *P=0.0275, Ordinary one-way ANOVA with Tukey post-hoc test, alpha = 0.05. B Fluorescent confocal image of Tau_441_ cells incubated for 24 h with either vehicle (V), 1 ×10^9^ P-EVs (P) or 1 ×10^9^ Tau_MBD_-EVs (Tau_MBD_) and stained for LysoTracker (in grey, N = nucleus). The histograms show, for the given treatments, LysoTracker-fluorescence analysed by flow-cytofluorimetry (mean ± s.e.m. of three biological replicates with ~2.7 ×10^3^ cells/condition, ****P<10^−15^, non-parametric Kruskal-Wallis one-way ANOVA, alpha = 0.05), the number and size of LysoTracker positive ELs analysed by confocal (mean ± s.e.m. of three biological replicates with 55, 54, 38 cells respectively, *P=0.0135, ***P=2 ×10^−4^, ****P<2.1 ×10^−5^, non-parametric Kruskal-Wallis one-way ANOVA, alpha = 0.05) C Tau_441_ cells incubated for 24 h with either vehicle (V), 1 ×10^9^ P-EVs (P) or 1 ×10^9^ Tau_MBD_-EVs (Tau_MBD_), stained for Hoechst and visulaized by fluorescence microscopy. The histograms show cell size, based on a Tau_441_ mask, and nuclear size, based on a hoechst mask, for the given treatments (mean ± s.d. of three biological replicates with 11.9 ×10^3^, 13.6 ×10^3^, 15.9 ×10^3^ cells/condition from three biological replicates). **P=0.0082, ****P<10^−15^, non-parametric Kruskal-Wallis one-way ANOVA, alpha = 0.05. D Cells with or without Tau_441_ expression (± induction) incubated for 24 h with either vehicle (V), 1 ×10^9^ P-EVs (P) or 1 ×10^9^ Tau_MBD_-EVs (Tau_MBD_). The histogram shows lactate dehydrogenase activity for the given treatments (mean ± s.d. of three biological replicates). ****P<1.5×10^−5^, ordinary two-way ANOVA with Tukey post-hoc test, alpha = 0.05.

Extracellular Tau_MBD_ seeds were shown to induce the aggregation of Tau in cultured C17.2 cells^6,36^ (Supplementary Fig 5A). To assess whether a similar phenotype may be induced by GFP-Tau_MBD_ transported by EVs, we incubated these latter on recipient cells engineered to express Tau_441_ fused to mCherry (here referred as Tau_441_, the largest splice variant of full-length Tau). Under these conditions, confocal microscopy revealed the induction of larger proportion of recipient cells carrying Tau-puncta when compared to cells treated in the absence (vehicle) or in the presence of EVs isolated from parental C17.2 cells (Fig 2D). Incubation with Tau_MBD_-EVs, when compared to parental-EVs, increased also the number, the size and the fluorescence intensity of Tau-puncta analysed at the single cell level (Fig 2E). Notably, analysis of Tau_441_-accumulation at the confocal microscope also revealed increased co-localisation with the ectopically expressed ELs marker GFP-CD63 as assessed by PCC determination (Fig 2F). This result was further substantiated by orthogonal reconstruction of stacked confocal planes demonstrating Tau_441_ accumulation within the lumen of GFP-CD63-positive organelles (Fig 2G).

**Figure 5.**
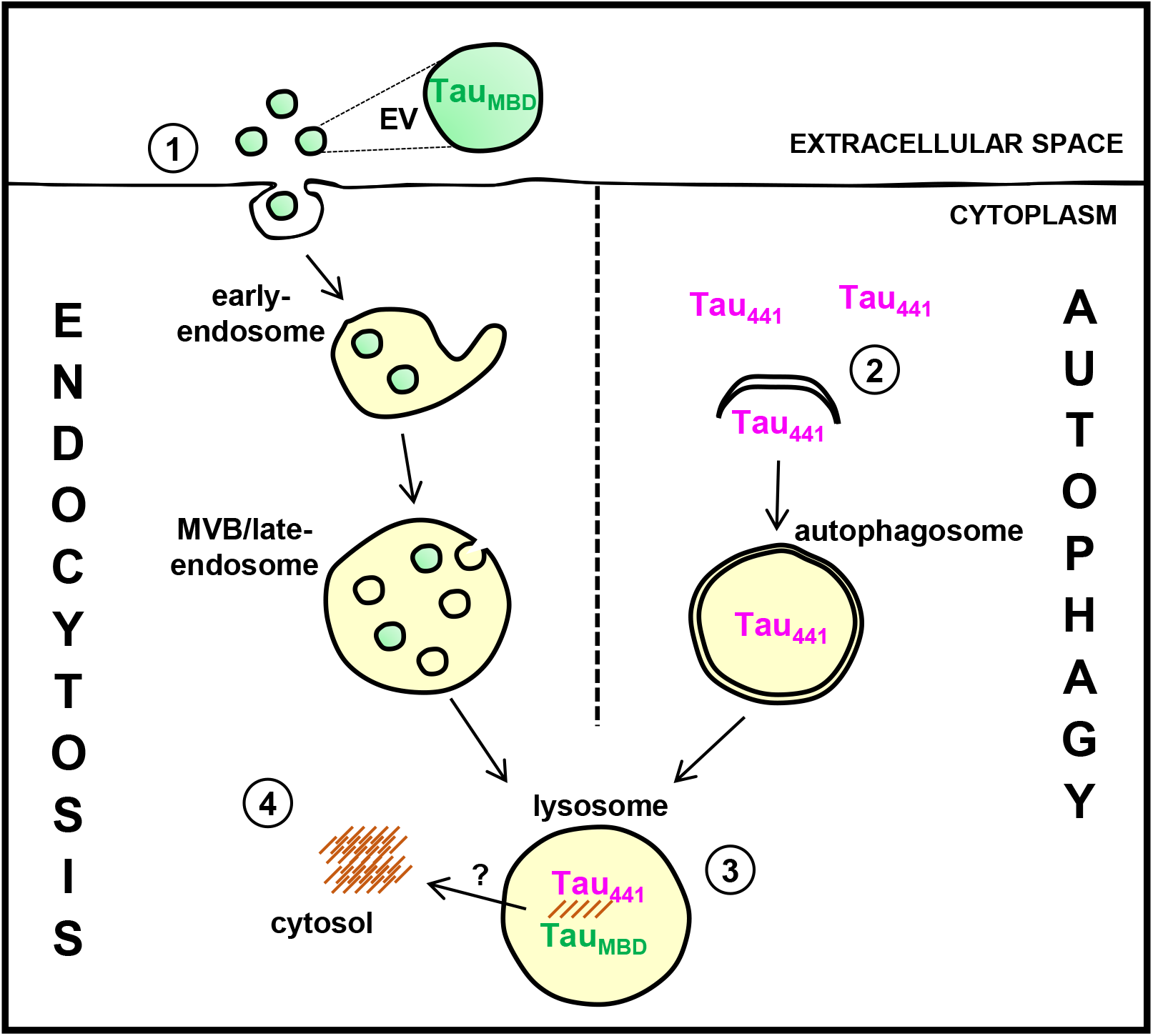
ELs as intracellular organelles allowing disease initiation. Scheme of intracellular and extracellular delivery of cargo to ELs. 1) Extracellular vesicles (EVs) transporting Tau_MBD_ are internalized by endocytosis in recipient cells, localize all along the endocytic pathway and accumulate in ELs. 2) Tau_441_ is a substrate of autophagy and it is delivered to ELs for degradation. 3) Extracellular Tau_MBD_ and intracellular Tau_441_ encounter, interact and propagate pathological conformations in ELs. 4) Eventual release into the cytosol as a consequence of excessive Tau accumulation.

### EVs mediated Tau_441_ accumulation in recipient cells requires autophagy

Tau_MBD_-EVs-induced accumulation of Tau_441_ in ELs hinted a likely participation of autophagy-mediated transport of cytosolic Tau_441_ to this acidic subcellular localisation, as previously reported^23^. In fact, nutrient-deprived Tau_441_-cells, which displayed an enhanced autophagy flux, acquired a Tau-puncta phenotype similar to that induced by Tau_MBD_-EVs in CD63-positive organelles (Supplementary Fig 5B and C). These data demonstrate that cytosolic Tau represents a substrate for autophagic degradation, which may become impaired by the presence of Tau_MBD_-EVs within ELs. Consistent with this hypothesis, Tau_MBD_-EVs induced the formation of Tau-puncta also in mouse embryonic fibroblasts (MEF) but not in MEF lacking ATG7 (ATG7-KO MEF), an E1-like activating enzyme indispensable for autophagosome assembly^43^ (Fig 2H). Lack of Tau-puncta in ATG7-KO MEF was not due to a defect in EVs-internalization since they displayed a Tau_MBD_-EVs internalization that was indistinguishable to that of normal MEF (Fig 2I).

### EVs transported TauMBD binds to recipient’s cells Tau_441_ in ELs

Co-localization of Tau_MBD_-EVs and Tau_441_ within ELs (Fig 3A) may favour their direct encounter. We studied previously protein-protein interaction and its subcellular localization utilizing a split GFP technology (biFC)^32,44^ based on a predominantly monomeric GFP^45^. Therefore, we isolated EVs from cells expressing GFP_1-10_ fused to Tau_MBD_ (GFP_1-10_-Tau_MBD_) and incubated them on cells expressing Tau_441_ fused to the remaining β-strand of GFP (T_11_-mCherry-Tau_441_) (Fig 3B) and analysed them by cytofluorimetry. This demonstrated the presence of a biFC signal in recipient cells (Fig 3C). The known sensitivity of GFP fluorescence to acidic conditions such as that expected in ELs may lead to an underestimation of the detected events. To assess the intracellular localization of biFC signal, we took advantage of the acidotropic fluorescent compound LysoTracker. To exclude a possible interference of LysoTracker with organelle acidification^46^, the compound was added just before the analysis and after controlling that the biFC signal was already present in the cells. Utilizing a slightly modified automated trained protocol^47^ to identify single LysoTracker-positive ELs, we established first that the mean biFC intensity in the ELs population was higher than that measured in the LysoTracker-negative cytoplasm (Fig 3D). Based on the observation that only a small portion of the ELs population was detected by biFC, we then analysed the biFC-positive ELs (arbitrarily defined as the 1% outliers) that most diverged from the mean biFC fluorescence derived from the whole ELs population. This allowed to establish that the biFC-positive ELs display Tau accumulation and increased size (Fig 3D). Overall, the data demonstrated the physical interaction between Tau_MBD_-EVs and Tau_441_ within ELs, which correlated with Tau_441_ accumulation in ELs presenting an enlarged size. This encounter of Tau species with distinct cellular origins in LAMP1-positive ELs led to the appearance of a conformational MC1-positive epitope, which reacts to abnormal conformations of Tau preceding Tau fibril formation and neurofibrillar toxicity^48^ as well as the double phosphorylation at Ser202/Thr305 recognized by the AT8 Tau antibody (Supplementary Fig 6).

The data described above indicate that the interaction between Tau_MBD_-EVs and Tau_441_ occurred preferentially within ELs and not in the cytosol. Next, we wanted to establish whether in our cell model an EVs-transported protein may reach the cytosol. For this, we selected the Cre recombinase, since the presence of this protein within the cell cytosol is detected with exquisite sensitivity with an adequate fluorescence reporter assay^15^. We first analysed a co-culture paradigm made of a stable cell line with inducible Cre expression and a cell line with an integrated loxP-dependent red-to-green reporter (Supplementary Fig 7A). Cytofluorimetric analysis established that recipient cells acquired green fluorescence indicative of successful transcellular transport of Cre produced by donor cells. However their relative number was not dependent on the level of Cre induction (Supplementary Fig 7B). Overall, the relative number of Cre-positive recipient cells was inferior to that of e.g. biFC positive cells even when the ratio of recipient to donor cells was increased 25:1 (Supplementary Fig 7C). When donor cells were transiently transfected with a Cre plasmid, this led to a cellular Cre expression higher than that obtained with the inducible system and the presence of detectable amounts of Cre in the EVs fraction (Supplementary Fig 7D). Incubation of these Cre-EVs on donor cells was equally inefficient in activating the red-to-green reporter and this did not change when the Cre-EVs treatment was performed in the presence of Tau_MBD_-EVs (Supplementary Fig 7E and F) although an eventual release into the cytosol as a consequence of excessive Tau accumulation cannot be excluded.

### EVs-mediated Tau_441_ accumulation causes lysosomal stress and cytotoxicity

Next, we investigated whether Tau_MBD_-EVs-mediated accumulation of Tau_441_ within ELs induced a potentially harmful cell response. With focus on lysosomal stress, we first analysed the nuclear translocation of a master transcription factor of lysosomal biogenesis (TFE3)^49,50^ in recipient cells and observed increased nuclear TFE3 localization in cells treated in the presence of Tau_MBD_-EVs when compared to vehicle or parental-EVs (Fig 4A). Under the same conditions, Tau_MBD_-EVs also increased number, dimension and fluorescence intensity of LysoTracker-positive ELs (Fig 4B). A significantly smaller but discernible effect was obtained also when cells were treated with parental EVs, i.e. without the Tau_MBD_-cargo (Fig 4B). Nevertheless, only the incubation with Tau_MBD_-EVs affected the overall cell morphology as observed in terms of reduced nuclear and cellular size (Fig 4C), a phenotype suggesting an ongoing cytotoxic process^51^. Consistent with this, an LDH-release assay demonstrated the induction of cytotoxicity in a manner that was dependent on the presence of Tau_MBD_-EVs as well as the induction of Tau_441_-expression (Fig 4D and Supplementary Fig 8).

## Discussion

EVs-mediated transport of macromolecules is a route of cell-to-cell communication with implications in health and disease. Whilst EVs biogenesis in donor cells is extensively investigated, the cellular mechanisms involved in EVs-cargo delivery in recipient cells needs further attention. For this reason, we focussed our investigations on the mechanisms of protein delivery mediated by EVs in the context of cell-to-cell propagation of Tau. In terms of a functionally relevant EVs-mediated protein transport, EVs-polypeptides present on the EVs-surface may directly interact with their respective receptors on the cell membrane. In contrast, for a luminal EVs-protein it is reasonable to assume that the cytosol may represent the main intracellular target site for the interaction with its effector. However, this latter would require a specific mechanism - for which experimental evidence is missing - that transfers the EVs-cargo to the cytosol. In our cell system, we confirmed that direct fusion of EVs at the plasma membrane is a possible but rather unlikely event of cargo delivery^15^. In contrast, and consistent with previous reports^18–20^, we observed a high rate of EVs internalization, possibly mimicking the internalization of e.g. high-density lipoproteins on their route to lysosomal degradation^52^. Cytosolic proteins are also delivered to lysosomes for degradation by a ubiquitous process accomplished by different forms of autophagy. Consistent with this model (Fig 5), we demonstrated herein that the initial encounter of EV-transported Tau_MBD_ with cellular Tau_441_ occurred in acidic organelles. For our experiments, we choose an EVs dose that was previously shown to be required for a functional activity in cultured cells^14^; and confirmed our results by co-culturing donor and recipient cells. Our data show that the encounter of a protein substrate of autophagy with an EVs-transported protein cargo resulted in a functionally relevant interaction between proteins originating from distinct cells. In particular, this observation attains a specific significance for a whole class of progressive proteinopathies, exemplified herein by tauopathies. Transcellular self-propagation of protein seeds with pathogenic conformations is proposed to explain the progressive nature of proteinopathies. For neurodegenerative disorders this prion-like mechanism appears to occur for most-neurodegeneration-linked proteins, including Aβ-peptides^53^, Tau^54^, huntingtin^55^, α-synuclein and TDP-43^56^. Extracellular seeds of these proteins cause the accumulation of normally functioning cognate protein in pathogenic conformations as shown e.g. by the cellular transfer of free-floating seeds of Tau^57^. EVs may act as vectors of prion infectivity^10,11^ and targeting of Tau in EVs is reported^58,59^. We now demonstrate that EVs-transported pathogenic Tau species induce a number of disease hallmarks in host cells, such as Tau accumulation, acquisition of a pro-pathologic epitope, ELs stress and cytotoxicity. Notably, our data are evidence that this adverse cascade of events may initiate within the acidic compartment of the host cell. Whether pathogenic Tau accumulated at this site ultimately is released in the cytosol or the specific location where Tau accumulated along the acidic degradative pathways are matter of future studies.

Turnover of cytosolic wild-type Tau, a long-lived protein, is mediated through the ubiquitin proteasome system and different forms of autophagy^60^. The age-dependent decline in the activity of these pathways is considered as the main cause for the build-up of aberrant protein forms; which in turns is possibly accelerated by the presence of protein aggregates^61–63^. Increased ELs number and aberrant ELs storage in neurons are hallmarks of neurodegeneration^64^. Our data showed that Tau_MBD_-EVs-induced Tau accumulation correlated with an ELs response characterized by morphological changes and the induction of TFE3 nuclear translocation. This molecular signature is also observed in the post-mortem human brain affected by neurodegeneration^65^. Autophagy appears upregulated in early AD, possibly as a response to protein accumulation, but then the autophagy flux - LC3, p62 and ELs enlargement – becomes progressively impaired^66–68^. Moreover, ELs markers are found enriched in brain protein deposits^69,70^. Considering the similarity with the observation made in our cell system, the effects reported herein may represent an underestimation of what may occur in non-dividing post-mitotic neurons.

Autophagy stimulation is generally considered as a viable solution to protect neurons from harmful protein aggregates^71^. However, delivering Tau to ELs for degradation may contribute to the formation of fibrillogenic Tau fragments^72^ and our data suggest that autophagy inhibition, rather than stimulation, may result in an approach to slow-down transcellular propagation of toxic Tau species. Since Tau is normally delivered to ELs^73,74^ (Fig 5, step 2), a possible mechanistic explanation for this hypothesis is supported by the finding that EVs-Tau_MBD_ binds and causes an aberrant Tau conformation ultimately impairing its degradation in ELs^75^ (Fig 5). Interestingly, it has been shown that within ELs Aβ-peptides^76^ or α-synuclein^77^ form aggregates and the acidic environment of ELs may favour fibril formation^78–81^. Based on our single ELs analysis, the existence of discrete subpopulations of ELs may be inferred each possibly specialized in the degradation of proteins with origin from one or more of the three main routes: the secretory, the autophagic or the endocytic pathways. The encounter of exogenous Tau_MBD_ seeds with cellular Tau demonstrated by the split GFP approach may thus occur in an acidic organelle that may take care of disposing extracellular and cytosolic proteins. Overall our data support the interplay of the endocytic and autophagic pathways in progressive proteinopathies.

## Acknowledgements

We thank A. Spang, E. Pecho-Vrieseling, members of L. Barile’s and M. Moretti’s laboratory and in particular all the members of our laboratory for discussions, critical advice and support during this study. PP is supported by grants from the Gelu Foundation and the Mecri Foundation.

## Author contributions

Conceptualization: PP and GP; Methodology: GP, MB, DM, SP, MM and PP; Investigation: GP, MB, DM and PP; Writing—original draft: PP; Writing—review & editing: GP, MB, DM, SP, MM and PP; Supervision: PP.

## Conflict of interest

The authors declare that they have no conflict of interest. PP and SP worked for AC Immune SA and own stock in the company. There are no patents, products in development, or marketed products to declare.

---

**Supplementary Figure 1 – DiD membranes are exchanged in co-cultures**

A 1D plots of DiD-fluorescence intensity for single DsRed cells co-cultured with parental cells (P) or DiD-labelled cells (DiD), the data are pooled from three flow-cytofluorimetry analyses (n ~5 ×10^3^ cells/replicate). The histogram shows the DiD fluorescence intensity determined for DsRed cells co-cultured as indicated (mean ± s.e.m. of three biological replicates with 35.2 ×10^3^, 54.1 ×10^3^ cells respectively). ****P<10^−15^, unpaired non-parametric Mann-Whitney test.

B Quantification of Fig 1B. The histogram shows the % of DsRed cells with DiO-fluorescence determined for DsRed cells co-cultured for 72 h with parental cells (P) or DiO-cells (DiO) (mean ± s.e.m. of six biological replicates with 48.4 ×10^3^, 56.5 ×10^3^ cells respectively). ****P=2.2 ×10^−3^, unpaired non-parametric Mann-Whitney test.

**Supplementary Figure 2 – Internalized DiD-EVs are localized in the endocytic pathway**

A Fluorescent confocal image of C17.2 cells transiently transfect with endocytic markers fused to mCherry (in magenta) incubated for 24 h with 1 ×10^9^ DiD-EVs (in green), co-emission of the two fluorophores results in a white colour and indicate localisation of DiD-EVs within endocytic positive organelles. Scale bar: 10 μm (1 μm for the enlarged inset).

B Fluorescent confocal image of a C17.2 cell incubated for 4 h with 11 ×0^9^ DiD-EVs and 10 μM of TRITC-dextran (DiD in green, dextran in magenta), co-emission of the two fluorophores results in a white colour and indicate localisation of DiD-EVs in dextran positive endocytic organelles. Scale bars: 10 μm (1 μm for the enlarged inset).

C 2D scatter plots of single cell DsRed and DiD fluorescence for three biological replicates (n ~1.4 ×10^3^ cells/replicate) by flow-cytofluorimetry for DsRed cells incubated with 1 ×10^9^ EVs (DiD-EVs) for 2 h in the absence (magenta) or presence of 33 μM dynasore (green) when compared to PBS control (vehicle, in grey).

**Supplementary Figure 3 – CD63-EVs are internalized in lamp1-positive ELs**

A 2D scatter plots of single cell DsRed and GFP fluorescence by flow-cytofluorimetry for the indicated cell populations for three biological replicates (n > 800 cells/condition). The histogram shows the GFP fluorescence intensity determined for DsRed cells co-cultured with parental cells (P) or GFP-CD63 expressing cells (CD63) (mean ± s.e.m. of four biological replicates with 1.5 ×10^3^, 1.0 ×10^3^ cells respectively). ****P<10^−15^, unpaired non-parametric Mann-Whitney test.

B Fluorescent confocal image for GFP-CD63 & DsRed co-culture (GFP in green, DsRed in magenta). The magnified image on the right includes an orthogonal reconstruction of 5 z-planes demonstrating internalized GFP-CD63 in a DsRed cell. Scale bar: 10 μm.

C Western blots of donor cell lysates (CL) and EVs lysates (EVs) for the EVs markers Alix and TSG101, or with a GFP-antibody detecting GFP-CD63 that is highly EVs enriched. Molecular weight markers are given on the left.

D 1D plots of single cell GFP-fluorescence intensity by flow-cytofluorimetry for parental cells (P cells) incubated with 1 ×10^9^ parental EVs (P-EVs, in black) or GFP-CD63-EVs (CD63, in green) for 24 h. The histogram shows GFP-fluorescence intensity for the given treatments (mean ± s.e.m. of three biological replicates with 24.6 ×10^3^, 24.4 ×10^3^ cells respectively), ****P<10^−15^, unpaired non-parametric Mann-Whitney test.

E Fluorescent confocal image of a C17.2 cell transiently transfect with mCherry-LAMP1 (in magenta) incubated for 24 h with 1 ×10^9^ CD63-EVs (in green), co-emission of the two fluorophores results in a white colour and indicate localisation of CD63-EVs within LAMP1 positive organelles. Scale bar: 10 μm (1 μm for the enlarged inset).

**Supplementary Figure 4 – EVs transported Tau_MBD_ protein is localized within the lumen of EVs**

A Scheme of the serial centrifugation procedure for enrichments of EVs from conditioned media of donor cells. P100 represents the pellet fraction recovered after a 100’000 x g centrifugation and S100 the respective supernatant.

B Representative experiment showing EVs concentration analysed by nanoparticle tracking analysis and protein concentration measured as absorbance at 280 nm in SEC fractions. The P100 and S100 recovered from the conditioned media of ~50 ×10^6^ C17.2 cells expressing GFP-Tau_MBD_ were applied to SEC columns separately. SEC fractions were pooled each three starting from F7 to F24.

C Assay to study luminal localization of Tau_MBD_ transported through EVs. 10 ×10^9^ GFP-Tau_MBD_-EVs were treated with either vehicle, 0.1% triton or 0.1 μg/mL trypsin and subsequently applied separately to a SEC column. After separation, fractions 7-to-15 (EVs-enriched) and fractions 16-to-24 were pooled. Western blots of EVs lysates for GFP-antibody detecting GFP-Tau_MBD_ and for the luminal EVs marker protein ALIX. Molecular weight markers are given on the left.

**Supplementary Figure 5 – Tau_441_ puncta induced by EVs transported Tau_MBD_ resemble those induced by Tau_MBD_ fibrils and starvation**

A Fluorescent confocal image of Tau_441_ cells (in grey, N = nucleus) incubated for 16 h with 0.5 μM of Tau_MBD_ fibrils. The image includes an orthogonal reconstruction of 14 focal z-planes demonstrating intracellular formation of Tau_441_ puncta. Scale bar: 10 μm.

B Tau_441_ cells treated with either vehicle (V), starved in HBSS (H) or starved in HBSS and simultaneously treated with 25 nM bafilomycin A1 (H & B) for 4 h. Western blots of Tau_441_ cell lysates for the autophagy marker LC3 and for housekeeping protein ACTIN. Molecular weight markers are given on the left. Fluorescent image of Tau_441_ cells incubated for 4 h with HBSS (in grey, N = nucleus). Scale bar: 10 μm. The histogram shows % of cells with a Tau-puncta phenotype for the given treatment (mean ± s.d. of three biological replicates with 1.6 ×10^3^, 1.3 ×10^3^ cells respectively). *P= 0.05, unpaired non-parametric Mann-Whitney test.

C Fluorescent confocal image of cells expressing Tau_441_ and GFP-CD63 (CD63) (Tau_441_ in magenta, CD63 in green, N = nucleus) incubated for 4 h with either vehicle or starved in HBSS, co-emission of the two fluorophores results in a white colour and indicate localisation of Tau_441_ within CD63 positive organelles. Scale bar: 10 μm (1 μm for the enlarged inset).

**Supplementary Figure 6 – EVs transported Tau_MBD_ induce the appearance of pathological Tau epitopes**

A Fluorescent confocal image of a T_11_-Tau_441_ cell (in magenta) incubated for 72 h with 6 ×10^9^ GFP-Tau_MBD_-EVs and stained either for MC1 or AT8 (in green), co-emission of the four fluorophores results in a white colour and indicate localisation of MC1/AT8 within and Tau_441_. Scale bar: 10 μm.

**Supplementary Figure 7 – EVs protein cargo transported through EVs rarely reach the cytosol of recipient cells**

A Scheme of co-culture assay to monitor trans-cytosolic transport of proteins using Cre-based recombination assay. Cre cells (in grey) co-cultured with DsRed cells (magenta) for 72 h results in recombined GFP cells (in green).

B Cre cells induced to express different amounts of Cre protein by tetracycline. Western blots of cell lysates for the Cre and for the housekeeping protein GAPDH. Molecular weight markers are given on the left. The histogram shows the % of DsRed cells with recombination in 72 h co-culture with Cre cells for the given concentration of tetracycline (mean ± s.d. of three biological replicates with 21.3 ×10^3^, 21.4 ×10^3^, 22.3 ×10^3^ cells respectively). Non-parametric Kruskal-Wallis one-way ANOVA, alpha = 0.05.

C Cre cells co-cultured with DsRed cells for 72 h at different Cre:DsRed ratios analysed by flow-cytofluorimetry. The histogram shows the % of DsRed cells with recombination for the given Cre (D):DsRed (R) ratio (mean ± s.d. of three biological replicates with 23.1 ×10^3^, 9.5 ×10^3^, 22.8 ×10^3^, 21.8 ×10^3^ cells respectively). **P= 6.7 ×10^3^, non-parametric Kruskal-Wallis one-way ANOVA, alpha = 0.05.

D WB analysis of donor cell lysates (CL) and isolated EVs fraction (EVs) for the indicated protein markers (Cre and TSG101). Cre protein expression were induced with the indicated concentrations of tetracycline or by transient transfection. Molecular weight markers are shown on the left of each panel.

E Confocal image showing a recipient DsRed cell (in magenta) that underwent to recombination (in green) when incubated for 72 h with 1 ×10^9^ Cre-EVs. Scale bar: 10 μm.

F DsRed cell incubated with 1-to-1 mixture of 1 ×10^9^ Cre-EVs with 1 ×10^9^ parental-EVs (P) or 1 ×10^9^ Cre-EVs with 1 ×10^9^ Tau_MBD_-EVs for 72 h analysed by flow-cytofluorimetry. The histogram shows the % of DsRed cells with recombination for the given treatment (mean ± s.d. of four biological replicates with 26.3 ×10^3^, 28.7 ×10^3^, 28.3 ×10^3^ cells respectively). **P= 4.7 ×10^3^, non-parametric Kruskal-Wallis one-way ANOVA, alpha = 0.05.

**Supplementary Figure 8 – Inducible Tau_441_ expression**

Fluorescent image of Tau_441_ inducible cells (in grey) incubated for 24 h with vehicle or 60 ng/mL of tetracycline and nuclei stained with Hoechst (in blue). Scale bar: 10 μm.

## References

1 Hardy, J. & Selkoe, D. J. The amyloid hypothesis of Alzheimer’s disease: progress and problems on the road to therapeutics. Science (New York, N.Y.) 297, 353–356, doi:10.1126/science.1072994 (2002).

2 Serrano-Pozo, A., Frosch, M. P., Masliah, E. & Hyman, B. T. Neuropathological alterations in Alzheimer disease. Cold Spring Harbor perspectives in medicine 1, a006189, doi:10.1101/cshperspect.a006189 (2011).

3 Braak, H. & Braak, E. Neuropathological stageing of Alzheimer-related changes. Acta neuropathologica 82, 239–259, doi:10.1007/bf00308809 (1991).

4 Freer, R. et al. A protein homeostasis signature in healthy brains recapitulates tissue vulnerability to Alzheimer’s disease. Science advances 2, e1600947, doi:10.1126/sciadv.1600947 (2016).

5 Clavaguera, F. et al. Transmission and spreading of tauopathy in transgenic mouse brain. Nature cell biology 11, 909–913, doi:10.1038/ncb1901 (2009).

6 Frost, B., Jacks, R. L. & Diamond, M. I. Propagation of tau misfolding from the outside to the inside of a cell. J Biol Chem 284, 12845–12852, doi:10.1074/jbc.M808759200 (2009).

7 Simons, M. & Raposo, G. Exosomes--vesicular carriers for intercellular communication. Current opinion in cell biology 21, 575–581, doi:10.1016/j.ceb.2009.03.007 (2009).

8 Kosaka, N. et al. Secretory mechanisms and intercellular transfer of microRNAs in living cells. J Biol Chem 285, 17442–17452, doi:10.1074/jbc.M110.107821 (2010).

9 Zhang, Y. et al. Secreted monocytic miR-150 enhances targeted endothelial cell migration. Molecular cell 39, 133–144, doi:10.1016/j.molcel.2010.06.010 (2010).

10 Alais, S. et al. Mouse neuroblastoma cells release prion infectivity associated with exosomal vesicles. Biology of the cell 100, 603–615, doi:10.1042/bc20080025 (2008).

11 Fevrier, B. et al. Cells release prions in association with exosomes. Proc Natl Acad Sci U S A 101, 9683–9688, doi:10.1073/pnas.0308413101 (2004).

12 Raposo, G. et al. B lymphocytes secrete antigen-presenting vesicles. The Journal of experimental medicine 183, 1161–1172, doi:10.1084/jem.183.3.1161 (1996).

13 Pegtel, D. M. et al. Functional delivery of viral miRNAs via exosomes. Proc Natl Acad Sci U S A 107, 6328–6333, doi:10.1073/pnas.0914843107 (2010).

14 Valadi, H. et al. Exosome-mediated transfer of mRNAs and microRNAs is a novel mechanism of genetic exchange between cells. Nature cell biology 9, 654–659, doi:10.1038/ncb1596 (2007).

15 Zomer, A. et al. In Vivo imaging reveals extracellular vesicle-mediated phenocopying of metastatic behavior. Cell 161, 1046–1057, doi:10.1016/j.cell.2015.04.042 (2015).

16 Skog, J. et al. Glioblastoma microvesicles transport RNA and proteins that promote tumour growth and provide diagnostic biomarkers. Nature cell biology 10, 1470–1476, doi:10.1038/ncb1800 (2008).

17 Parolini, I. et al. Microenvironmental pH is a key factor for exosome traffic in tumor cells. J Biol Chem 284, 34211–34222, doi:10.1074/jbc.M109.041152 (2009).

18 Costa Verdera, H., Gitz-Francois, J. J., Schiffelers, R. M. & Vader, P. Cellular uptake of extracellular vesicles is mediated by clathrin-independent endocytosis and macropinocytosis. Journal of controlled release : official journal of the Controlled Release Society 266, 100–108, doi:10.1016/j.jconrel.2017.09.019 (2017).

19 Mulcahy, L. A., Pink, R. C. & Carter, D. R. Routes and mechanisms of extracellular vesicle uptake. Journal of extracellular vesicles 3, doi:10.3402/jev.v3.24641 (2014).

20 Heusermann, W. et al. Exosomes surf on filopodia to enter cells at endocytic hot spots, traffic within endosomes, and are targeted to the ER. The Journal of cell biology 213, 173–184, doi:10.1083/jcb.201506084 (2016).

21 Settembre, C., Fraldi, A., Medina, D. L. & Ballabio, A. Signals from the lysosome: a control centre for cellular clearance and energy metabolism. Nature reviews. Molecular cell biology 14, 283–296, doi:10.1038/nrm3565 (2013).

22 Lee, M. J., Lee, J. H. & Rubinsztein, D. C. Tau degradation: the ubiquitin-proteasome system versus the autophagy-lysosome system. Progress in neurobiology 105, 49–59, doi:10.1016/j.pneurobio.2013.03.001 (2013).

23 Wang, Y. & Mandelkow, E. Degradation of tau protein by autophagy and proteasomal pathways. Biochemical Society transactions 40, 644–652, doi:10.1042/bst20120071 (2012).

24 Nikoletopoulou, V., Papandreou, M. E. & Tavernarakis, N. Autophagy in the physiology and pathology of the central nervous system. Cell death and differentiation 22, 398–407, doi:10.1038/cdd.2014.204 (2015).

25 Yang, Z. & Klionsky, D. J. Mammalian autophagy: core molecular machinery and signaling regulation. Current opinion in cell biology 22, 124–131, doi:10.1016/j.ceb.2009.11.014 (2010).

26 Malik, B. R., Maddison, D. C., Smith, G. A. & Peters, O. M. Autophagic and endo-lysosomal dysfunction in neurodegenerative disease. Molecular brain 12, 100, doi:10.1186/s13041-019-0504-x (2019).

27 Wang, C., Telpoukhovskaia, M. A., Bahr, B. A., Chen, X. & Gan, L. Endo-lysosomal dysfunction: a converging mechanism in neurodegenerative diseases. Current opinion in neurobiology 48, 52–58, doi:10.1016/j.conb.2017.09.005 (2018).

28 Rost, B. R. et al. Optogenetic acidification of synaptic vesicles and lysosomes. Nature neuroscience 18, 1845–1852, doi:10.1038/nn.4161 (2015).

29 Van Engelenburg, S. B. & Palmer, A. E. Imaging type-III secretion reveals dynamics and spatial segregation of Salmonella effectors. Nature methods 7, 325–330, doi:10.1038/nmeth.1437 (2010).

30 Friedman, J. R., Webster, B. M., Mastronarde, D. N., Verhey, K. J. & Voeltz, G. K. ER sliding dynamics and ER-mitochondrial contacts occur on acetylated microtubules. The Journal of cell biology 190, 363–375, doi:10.1083/jcb.200911024 (2010).

31 Rowland, A. A., Chitwood, P. J., Phillips, M. J. & Voeltz, G. K. ER contact sites define the position and timing of endosome fission. Cell 159, 1027–1041, doi:10.1016/j.cell.2014.10.023 (2014).

32 Foglieni, C. et al. Split GFP technologies to structurally characterize and quantify functional biomolecular interactions of FTD-related proteins. Scientific reports 7, 14013, doi:10.1038/s41598-017-14459-w (2017).

33 Thery, C., Amigorena, S., Raposo, G. & Clayton, A. Isolation and characterization of exosomes from cell culture supernatants and biological fluids. Current protocols in cell biology Chapter 3, Unit 3.22, doi:10.1002/0471143030.cb0322s30 (2006).

34 Schrader-Fischer, G. & Paganetti, P. A. Effect of alkalizing agents on the processing of the beta-amyloid precursor protein. Brain research 716, 91–100, doi:10.1016/0006-8993(96)00002-9 (1996).

35 Ryder, E. F., Snyder, E. Y. & Cepko, C. L. Establishment and characterization of multipotent neural cell lines using retrovirus vector-mediated oncogene transfer. Journal of Neurobiology 21, 356–375, doi:10.1002/neu.480210209 (1990).

36 Holmes, B. B. et al. Heparan sulfate proteoglycans mediate internalization and propagation of specific proteopathic seeds. Proc Natl Acad Sci U S A 110, E3138–E3147, doi:10.1073/pnas.1301440110 (2013).

37 Gardiner, C. et al. Techniques used for the isolation and characterization of extracellular vesicles: results of a worldwide survey. Journal of extracellular vesicles 5, 32945, doi:10.3402/jev.v5.32945 (2016).

38 Thery, C. et al. Minimal information for studies of extracellular vesicles 2018 (MISEV2018): a position statement of the International Society for Extracellular Vesicles and update of the MISEV2014 guidelines. Journal of extracellular vesicles 7, 1535750, doi:10.1080/20013078.2018.1535750 (2018).

39 Macia, E. et al. Dynasore, a cell-permeable inhibitor of dynamin. Developmental cell 10, 839–850, doi:10.1016/j.devcel.2006.04.002 (2006).

40 Miesenbock, G., De Angelis, D. A. & Rothman, J. E. Visualizing secretion and synaptic transmission with pH-sensitive green fluorescent proteins. Nature 394, 192–195, doi:10.1038/28190 (1998).

41 Vogt, S., Stadlmayr, G., Grillari, J., Rüker, F. & Wozniak-Knopp, G. in Current Topics in Biochemical Engineering (ed IntechOpen) (2019).

42 Andreu, Z. & Yanez-Mo, M. Tetraspanins in extracellular vesicle formation and function. Frontiers in immunology 5, 442, doi:10.3389/fimmu.2014.00442 (2014).

43 Komatsu, M. et al. Impairment of starvation-induced and constitutive autophagy in Atg7-deficient mice. The Journal of cell biology 169, 425–434, doi:10.1083/jcb.200412022 (2005).

44 Ulrich, G. et al. Phosphorylation of nuclear Tau is modulated by distinct cellular pathways. Scientific reports 8, 17702, doi:10.1038/s41598-018-36374-4 (2018).

45 Cabantous, S., Terwilliger, T. C. & Waldo, G. S. Protein tagging and detection with engineered self-assembling fragments of green fluorescent protein. Nature biotechnology 23, 102–107, doi:10.1038/nbt1044 (2005).

46 Probes, M. Molecular Probes Handbook A Guide to Fluorescent Probes and Labeling Technologies(2010).

47 Morone, D., Marazza, A., Bergmann, T. J. & Molinari, M. Deep learning approach for quantification of organelles and misfolded polypeptides delivery within degradative compartments. Molecular Biology of the Cell 0, mbc.E20-04-0269, doi:10.1091/mbc.E20-04-0269.

48 Jicha, G. A., Berenfeld, B. & Davies, P. Sequence requirements for formation of conformational variants of tau similar to those found in Alzheimer’s disease. Journal of neuroscience research 55, 713–723, doi:10.1002/(sici)1097-4547(19990315)55:6<713::aid-jnr6>3.0.co;2-g (1999).

49 Martina, J. A. et al. The Nutrient-Responsive Transcription Factor TFE3 Promotes Autophagy, Lysosomal Biogenesis, and Clearance of Cellular Debris. Science Signaling 7, ra9, doi:10.1126/scisignal.2004754 (2014).

50 Settembre, C. et al. TFEB links autophagy to lysosomal biogenesis. Science (New York, N.Y.) 332, 1429–1433, doi:10.1126/science.1204592 (2011).

51 Cummings, B. S. & Schnellmann, R. G. Measurement of cell death in mammalian cells. Current protocols in pharmacology Chapter 12, Unit 12.18, doi:10.1002/0471141755.ph1208s25 (2004).

52 Miller, N. E., Weinstein, D. B. & Steinberg, D. Binding, internalization, and degradation of high density lipoprotein by cultured normal human fibroblasts. Journal of lipid research 18, 438–450 (1977).

53 Eisele, Y. S. et al. Peripherally applied Abeta-containing inoculates induce cerebral beta-amyloidosis. Science (New York, N.Y.) 330, 980–982, doi:10.1126/science.1194516 (2010).

54 Clavaguera, F., Duyckaerts, C. & Haik, S. Prion-like properties of Tau assemblies. Current opinion in neurobiology 61, 49–57, doi:10.1016/j.conb.2019.11.022 (2020).

55 Pecho-Vrieseling, E. et al. Transneuronal propagation of mutant huntingtin contributes to non-cell autonomous pathology in neurons. Nature neuroscience 17, 1064–1072, doi:10.1038/nn.3761 (2014).

56 Peled, S. et al. Single cell imaging and quantification of TDP-43 and alpha-synuclein intercellular propagation. Scientific reports 7, 544, doi:10.1038/s41598-017-00657-z (2017).

57 Katsinelos, T. et al. Unconventional Secretion Mediates the Trans-cellular Spreading of Tau. Cell reports 23, 2039–2055, doi:10.1016/j.celrep.2018.04.056 (2018).

58 Pérez, M., Avila, J. & Hernández, F. Propagation of Tau via Extracellular Vesicles. Front Neurosci 13, 698–698, doi:10.3389/fnins.2019.00698 (2019).

59 Wang, Y. et al. The release and trans-synaptic transmission of Tau via exosomes. Molecular neurodegeneration 12, 5–5, doi:10.1186/s13024-016-0143-y (2017).

60 Caballero, B. et al. Interplay of pathogenic forms of human tau with different autophagic pathways. Aging Cell 17, e12692, doi:10.1111/acel.12692 (2018).

61 Menzies, F. M., Fleming, A. & Rubinsztein, D. C. Compromised autophagy and neurodegenerative diseases. Nature reviews. Neuroscience 16, 345–357, doi:10.1038/nrn3961 (2015).

62 Nixon, R. A. The role of autophagy in neurodegenerative disease. Nature medicine 19, 983–997, doi:10.1038/nm.3232 (2013).

63 Chen, X. et al. Promoting tau secretion and propagation by hyperactive p300/CBP via autophagy-lysosomal pathway in tauopathy. Molecular neurodegeneration 15, 2, doi:10.1186/s13024-019-0354-0 (2020).

64 Nixon, R. A. et al. Extensive involvement of autophagy in Alzheimer disease: an immuno-electron microscopy study. Journal of neuropathology and experimental neurology 64, 113–122, doi:10.1093/jnen/64.2.113 (2005).

65 Martini-Stoica, H., Xu, Y., Ballabio, A. & Zheng, H. The Autophagy-Lysosomal Pathway in Neurodegeneration: A TFEB Perspective. Trends in neurosciences 39, 221–234, doi:10.1016/j.tins.2016.02.002 (2016).

66 Bordi, M. et al. Autophagy flux in CA1 neurons of Alzheimer hippocampus: Increased induction overburdens failing lysosomes to propel neuritic dystrophy. Autophagy 12, 2467–2483, doi:10.1080/15548627.2016.1239003 (2016).

67 Piras, A., Collin, L., Grüninger, F., Graff, C. & Rönnbäck, A. Autophagic and lysosomal defects in human tauopathies: analysis of post-mortem brain from patients with familial Alzheimer disease, corticobasal degeneration and progressive supranuclear palsy. Acta Neuropathologica Communications 4, 22, doi:10.1186/s40478-016-0292-9 (2016).

68 Wong, E. & Cuervo, A. M. Autophagy gone awry in neurodegenerative diseases. Nature neuroscience 13, 805–811, doi:10.1038/nn.2575 (2010).

69 Gowrishankar, S. et al. Massive accumulation of luminal protease-deficient axonal lysosomes at Alzheimer’s disease amyloid plaques. Proceedings of the National Academy of Sciences, 201510329, doi:10.1073/pnas.1510329112 (2015).

70 Hassiotis, S. et al. Lysosomal LAMP1 immunoreactivity exists in both diffuse and neuritic amyloid plaques in the human hippocampus. The European journal of neuroscience 47, 1043–1053, doi:10.1111/ejn.13913 (2018).

71 Yuan, Z., Yidan, Z., Jian, Z., Xiangjian, Z. & Guofeng, Y. Molecular mechanism of Autophagy: Its role in the therapy of Alzheimer’s disease. Current neuropharmacology, doi:10.2174/1570159x18666200114163636 (2020).

72 Wang, Y. et al. Tau fragmentation, aggregation and clearance: the dual role of lysosomal processing. Human molecular genetics 18, 4153–4170, doi:10.1093/hmg/ddp367 (2009).

73 Ji, C., Tang, M., Zeidler, C., Hohfeld, J. & Johnson, G. V. BAG3 and SYNPO (synaptopodin) facilitate phospho-MAPT/Tau degradation via autophagy in neuronal processes. Autophagy 15, 1199–1213, doi:10.1080/15548627.2019.1580096 (2019).

74 Vaz-Silva, J. et al. Endolysosomal degradation of Tau and its role in glucocorticoid-driven hippocampal malfunction. The EMBO journal 37, doi:10.15252/embj.201899084 (2018).

75 Guix, F. X. The interplay between aging-associated loss of protein homeostasis and extracellular vesicles in neurodegeneration. Journal of neuroscience research 98, 262–283, doi:10.1002/jnr.24526 (2020).

76 Brewer, G. J. et al. Age-Related Intraneuronal Aggregation of Amyloid-beta in Endosomes, Mitochondria, Autophagosomes, and Lysosomes. Journal of Alzheimer’s disease : JAD 73, 229–246, doi:10.3233/jad-190835 (2020).

77 Tsujimura, A. et al. Lysosomal enzyme cathepsin B enhances the aggregate forming activity of exogenous α-synuclein fibrils. Neurobiology of disease 73, 244–253, doi:10.1016/j.nbd.2014.10.011 (2015).

78 McGlinchey, R. P., Jiang, Z. & Lee, J. C. Molecular origin of pH-dependent fibril formation of a functional amyloid. Chembiochem : a European journal of chemical biology 15, 1569–1572, doi:10.1002/cbic.201402074 (2014).

79 Moriarty, G. M. et al. A pH-dependent switch promotes beta-synuclein fibril formation via glutamate residues. The Journal of biological chemistry 292, 16368–16379, doi:10.1074/jbc.M117.780528 (2017).

80 Pfefferkorn, C. M., McGlinchey, R. P. & Lee, J. C. Effects of pH on aggregation kinetics of the repeat domain of a functional amyloid, Pmel17. Proceedings of the National Academy of Sciences 107, 21447, doi:10.1073/pnas.1006424107 (2010).

81 Uversky, V. N., Li, J. & Fink, A. L. Evidence for a partially folded intermediate in alpha-synuclein fibril formation. The Journal of biological chemistry 276, 10737–10744, doi:10.1074/jbc.M010907200 (2001).

